# Emigration Effects on Estimates of Age- and Sex-specific Survival of Small Mammals

**DOI:** 10.1101/2021.08.07.455530

**Authors:** Matthew J. Weldy, Damon B. Lesmeister, Clinton W. Epps

**Affiliations:** Pacific Northwest Research Station, U.S.D.A. Forest Service, Corvallis, Oregon, USA; Department of Fisheries and Wildlife, Oregon State University, Corvallis, Oregon, USA

**Keywords:** Apparent survival, emigration, immigration, Sciuridae, site-fidelity, vital rates

## Abstract

1. Age- and sex-specific survival estimates are crucial to understanding important life-history characteristics and variation in these estimates can be a key driver of population dynamics. When estimating survival using Cormack–Jolly–Seber (CJS) models and capture–recapture data, emigration is typically assumed to have a negligible effect on estimates such that apparent survival is indistinguishable from true survival. Consequently, especially for populations or age classes with high dispersal rates, apparent survival estimates are often biased low and temporal patterns in survival might be masked when site fidelity varies temporally.
2. We used 9 years of annual mark-recapture data to estimate age-, sex-, and time-specific apparent survival of Humboldt’s flying squirrels (*Glaucomys oregonensis*) and Townsend’s chipmunks (*Neotamias townsendii*). For Humboldt’s flying squirrels, these estimates support a small body of research investigating potential variation of survival among age and sex classes, but age- and sex-specific survival has not been evaluated for Townsend’s chipmunks. We also quantified the effects of age- and sex-specific emigration on confounded estimates of apparent survival.
3. Our estimates of juvenile flying squirrel survival were high relative to other small mammal species and estimates for both species were variable among years. We found survival differed moderately among age and sex classes for Humboldt’s flying squirrels, but little among age and sex classes for Townsend’s chipmunks, and that the degree to which emigration confounded apparent survival estimates varied substantially among years. Without correcting for emigration, apparent survival estimates were lower and temporal variation was obscured, particularly for male Humboldt’s flying squirrels and female Townsend’s chipmunks.
4. Our results demonstrate that emigration can influence commonly used estimates of apparent survival. Unadjusted estimates confounded the interpretation of differences in survival between age and sex classes and masked potential temporal patterns in survival because the magnitude of adjustment varied among years. We conclude that apparent survival estimators are robust during some time periods; however, when emigration rates vary in time the effects of emigration should be carefully considered and accounted for, especially in comparative studies and those with policy and conservation implications.

## 1 Introduction

Variation in survival rates can be a key driver of population dynamics (Cole, 1954), and thus is vital for the study of population demography and life history (Franklin et al., 1996). Within a population, the relative survival of juveniles and adults (Charlesworth, 1994) or males and females (Promislow, 1992) can help inform which factors are important to population dynamics and how populations will change through time (Morrison & Hik, 2007). In mammals, adult and juvenile survival are often correlated, although juvenile survival is usually lower and more variable (Promislow & Harvey, 1990). This variation is thought to be regulated by juvenile life history characteristics such as natal dispersal (Rödel et al., 2015), or increased sensitivity to limited food (Jackson et al., 2001), thermoregulatory stress (Rödel et al., 2004), and predation (Garrett & Franklin, 1988). Similarly, there is a long-standing belief that males exhibit lower survival rates than females (Vinogradov, 1998) because of exposure to higher costs of dispersal associated with locating and competing for mates (Promislow, 2003). Despite their importance in understanding the relative influence of survival on population dynamics, age-, sex-, and time-specific estimates of survival are unavailable for many small mammal species because obtaining suitable data is challenging.

One common method for estimating survival is the Cormack–Jolly–Seber (CJS) model, which jointly estimates apparent survival and recapture probabilities (Cormack, 1964; Jolly, 1965; Seber, 1965). Apparent survival estimates from capture-recapture data and CJS models are commonly interpreted as estimates of survival; however, the estimated parameter is the product of true survival and site-fidelity (Lebreton et al., 1992) because individual survival is indistinguishable from permanent emigration. If emigration is permanent or non-random, CJS survival probability estimates will be biased low (Schaub et al., 2004) and the magnitude of bias can be large (e.g., Cooper et al., 2008; Horton & Letcher, 2008). A number of approaches have been suggested to deal with this bias. For example, a multistate approach allows for separation of movement and survival probabilities (Brownie et al., 1993), the robust design approach (Pollock et al., 1990) can account for temporary emigration, and data integrations can allow joint estimation of true survival and site fidelity (e.g., Burnham, 1993). More recently, Gilroy et al. (2012) and Schaub and Royle (2014) developed CJS model extensions that adjusted estimates of apparent survival with those of site fidelity. In some cases, estimates of apparent survival will suffice for conservation or management. However, when little is known about species-specific variation in survival (age or sex variation) or patterns of dispersal, inferences based on apparent survival estimates could mask important spatial or temporal variation in true survival.

Forest-adapted small mammals are important to forest heath as prey species and dispersal agents of hypogeous fungi and spermatophyte seeds (Trappe et al., 2009); yet, few papers have estimated movement rates (emigration, immigration, or site-fidelity) for these species, and thus unbiased estimates of survival are rare or non-existent. We focused our analyses on two small mammal species, Humboldt’s flying squirrels (*Glaucomys oregonensis*; hereafter flying squirrel) and Townsend’s chipmunks (*Neotamias townsendii*; hereafter chipmunk). Flying squirrels have been characterized as a potentially K-selected species, with survival that is higher than similar-sized mammals (Smith, 2007; Villa et al., 1999) and varies little across time (Lehmkuhl et al., 2006). However, other demographic characteristics such as abundance (Weldy et al. 2019), sex ratio (Rosenberg & Anthony, 1992), and recruitment (Weldy et al. 2020) vary substantially across time. Much less is known about chipmunk demography, but they have been characterized as an r-selected species with population growth rates primarily driven by recruitment and generally lower survival which can exhibit substantial temporal variation (Weldy et al., 2020). Little is known about variation in survival among age or sex classes for either species, but differential juvenile mortality rates (Forsman et al., 2004) or sex-specific effects could cause negative biases in apparent survival estimates.

Our objectives for this study were to estimate age-, sex-, and time-specific annual survival and recapture probability for flying squirrels and chipmunks captured in old forests during a relatively undisturbed 9-year period. We used mark-recapture data and two estimators to quantify sensitivity of survival estimates to variation in movement probabilities: 1) a CJS estimator which jointly estimates apparent survival and recapture probability, and 2) an integrated modelling approach to estimate immigration rates, which we used to derive site-fidelity rates and emigration-adjusted survival. We hypothesized that survival would vary among age and sex classes and that differences in apparent survival among age and sex classes was confounded by variation in emigration rates. For juveniles of both species, we predicted lower survival probabilities and higher immigration rates relative to subadults and adults, but that adjusting apparent survival for emigration would reduce differences in survival among age-classes (Dobson, 1982). We also predicted that males of both species would have lower survival and higher immigration rates relative to females because male mammals typically disperse more frequently and farther, and have higher predation risk and resource acquisition costs (Lemaître et al., 2020).

## 2 Materials and Methods

### 2.1 Study Area

We collected field data annually during September–November 2011–2019 on nine sites in the H. J. Andrews Experimental Forest on the west slope of the Cascade Mountains in Oregon, United States (44°14’ N, 122°10’ W). The study sites were all located in a late-successional forest (> 400 years) dominated by large Douglas-fir (*Pseudotsuga menziesii*), western hemlock (*Tsuga heterophylla*), and Pacific silver fir (*Abies amabilis*; Schulze & Lienkaember, 2015). Understory characteristics on the study sites ranged from open understories to dense shrubs and common understory vegetation included black berry, raspberry, and salmonberry (*Rubus* spp.), common snowberry (*Symphoricarpos albus*), deer fern (*Blechnum spicant*), huckleberry (*Vaccinium* spp.), Oregon grape (*Mahonia aquifolium*), oxalis (*Oxalis* spp.), salal (*Gaultheria shallon*), sword fern (*Polystichum munitum*), and vine maple (*Acer circinatum*).

Weather on the study sites was typically warm and dry from May–September, and cool and wet from October–April when approximately 80% of the annual precipitation occurs. Annual precipitation primarily consists of rain at elevations < 1,000 m and snow at elevations ≥ 1,000 m (Bierlmaler & McKee, 1989). At 605 m, 30-year (1981–2010) averages were 1,955.8 mm of precipitation, 4.3° C minimum temperature, and 15.6° C maximum temperature (PRISM Climate Group, 2004). Monthly temperature and precipitation varied seasonally and inter-annually during the period of our study (Figure S1).

### 2.2 Data Collection

The nine study sites (7.84 ha each) were randomly selected across gradients of elevation (range = 683–1,244 m) and canopy openness (range = 0–40%). The average distance among sites was 2,963 m (range = 1,078–5,940 m). As described by Weldy et al. (2020), at each site we established and conducted live-trapping at 64 stations, each with two traps, arranged in an 8 × 8 array with 40 m between stations. For each animal, we recorded age, body mass (g), reproductive condition, species, and sex (Villa et al. 1991; Gashwiler, 1976). Live-trapping protocols were approved by the Oregon State University’s Institutional Animal Care and Use Committee (ACUP #4191 2011–2013; #4590 2014–2016; #4959 2017–2019) and were consistent with the American Society of Mammalogists guidelines for the use of wild mammals in research and education (Sikes et al., 2016).

### 2.3 Analytical Methods

We estimated apparent annual survival (***φ***) and recapture probability (*p*) for flying squirrels and chipmunks using mark-recapture data and CJS models (Cormack, 1964; Jolly, 1965; Seber, 1965) in a state-space formulation (Gimenez et al., 2007), with an additional sub model to estimate immigration rates (*I*). This hierarchical model consists of two state processes and one observation process. The first state process, ***φ*** (the probability of surviving and remaining in the study area), was linked with the observation processes *p* (the capture probability of a marked individual; Lebreton et al., 1992). The second state process, *I*, was independent from the ***φ*** process and the *p* process and was used to derive site-fidelity rates (*r*) and emigration-adjusted survival estimates (***φ***_adjusted_).

For the ***φ*** process, we first defined a latent variable *z_i,t_* as the true state of individual *i* at time *t*, where a value of 1 indicated *i* was alive at *t* and a value of 0 indicated *i* was dead at *t*. We also defined a vector ***f***, where *f_i_* was the first capture occasion for individual *i*. We modelled the probability that *i* was alive at *t*+1, conditional on first capture and being alive at *t*, as a Bernoulli trial where the success probability is the product of ***φ**_a,s,t_* and *z_i,t_*

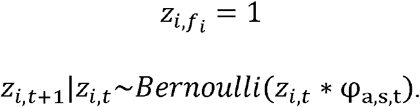

We modelled variation in ***φ*** as a logit-linear function of an age-specific (*a*) intercept for *i* at *t*, where ages ranged 1–3 (juvenile, subadult, adult) for flying squirrels and 1–2 (juvenile, adult) for chipmunks, and an additive age- and sex-specific (*s*) effect, where the effect of sex differed by age, the sex variable was defined as 0 for males and 1 for females, and we included a zero-centered age-, sex-, and time-specific random effect with standard deviation ***σ***_a,s,t_.

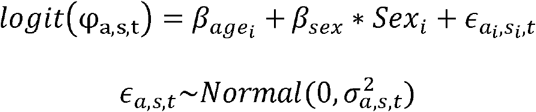

We used a standard observation process for capture-recapture data *y_i,t_*, where recaptures at each occasion from the second to the last trapping occasion were modelled as Bernoulli trials with success probability *p_i,t_*. To determine the most supported model structure for the observation process we considered nine logit-linear model structures (seven univariate, two bivariate) for both species to account for variation in *p* (Table 1).

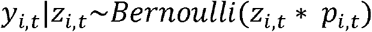

**TABLE 1.**
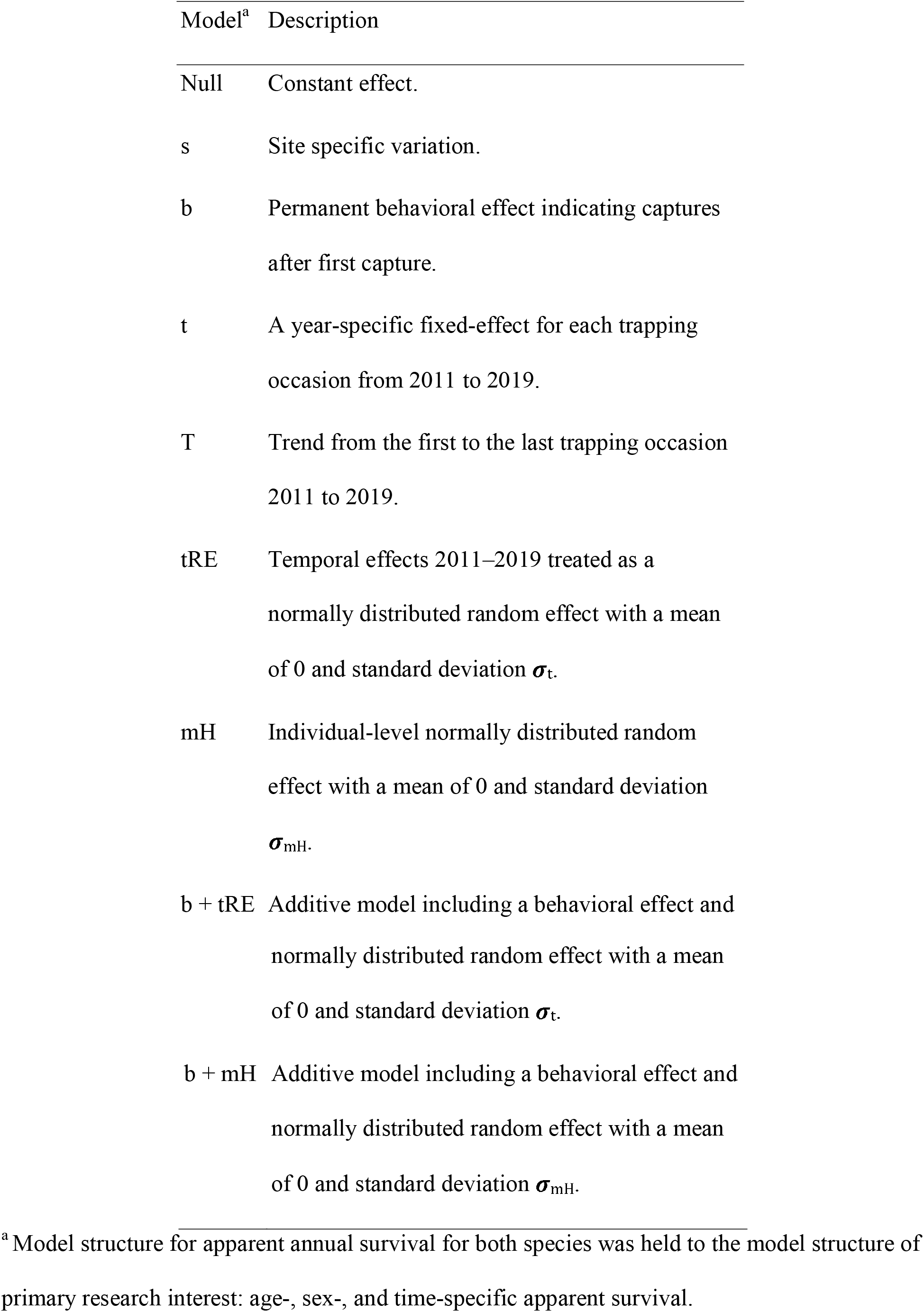
Description of variables considered in Cormack–Jolly–Seber models of recapture probability (*p*) for Humboldt’s flying squirrels (*Glaucomys oregonensis*) and Townsend’s chipmunks (*Neotamias townsendii*) fitted using mark-recapture data, 2011–2019, recorded in the H. J. Andrews Experimental Forest, near Blue River, Oregon.

We used Poisson regression to estimate age-, sex, and time-specific immigration rates (*I_a,s,t_*) during *t* from *t* = 2 to the number of occasions. The response variable (*Imm_a,s,t_*) was the age-, sex-, and time-specific counts of captured unmarked individuals.

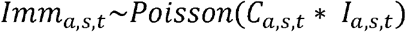

We modelled variation in *I_a,s,t_* as a log-linear function of an age-specific effect, a sex-specific effect, and a zero-centered normally distributed time-specific random effect with standard deviation ***σ**_t_*.

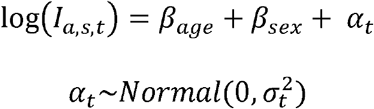

We assumed that age- and sex-specific emigration at *t* was equal to *I_a,s,t+1_*, and that age- and sex-specific *r_a,s,t_* was the complement to emigration (i.e., *r_a,s,t_* = 1 – emigration_*a,s,t*_). These assumptions were reasonable on the study sites, which were randomly placed within a large, continuous, old late-successional forest. We then derived estimates of emigration-adjusted survival (***φ***_adjusted_), defined as:

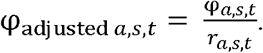

For the observation process, we used the Watanabe-Akaike information criterion (WAIC; Watanabe, 2010) to select the most supported model structure for *p* (Hooten & Hobbs, 2015; Vehtari et al., 2017). We considered the model with the smallest WAIC value and highest model support weight (ω) the most supported model. We used the relative change in WAIC (ΔWAIC) to evaluate models relative to the top-ranking model, and because the estimate of WAIC is sensitive to the sample distribution, we estimated the 95% credible interval (CI) for the difference in WAIC from the top-ranking model. We assessed the meaningfulness of a difference between two models based on the degree to which the 95% CI for the difference did or did not overlap zero.

We evaluated goodness-of-fit for the CJS model using a posterior predictive check approach (Gelman et al., 2013) to estimate a Bayesian p-value (Meng, 1994). The data are binary and standard fit statistics are uninformative about model fit. Thus, a Bayesian *p*-value was derived as the proportion of times that Chi-squared test statistics (Pearson, 1900) calculated for simulated datasets were higher than chi-squared test statistics for an aggregation of the observed datasets (i.e., individual row sums; Royle et al., 2014). Perfect agreement between the observed and simulated datasets occurs when the Bayesian *p*-value equals 0.5.

We conducted all analyses using R version 3.6.1 (R Core Team, 2020). The models were fitted using JAGS software version 4.3.0 (Plummer, 2003) through the R2jags package version 0.6-1 (Su & Yajima, 2009). We used diffuse priors for all parameters and evaluated prior sensitivity using two sets of priors. During model selection steps, each model was estimated with three independent chains of 5,000 iterations following a burn-in period of 2,000 iterations. For inference, the top-ranking models for flying squirrels and chipmunks were estimated with three independent 50,000 iteration chains each following a burn-in period of 50,000 iterations. We assessed model convergence by visual examination of trace plots and computed the Brooks–Gelman–Rubin convergence diagnostic (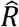; Brooks & Gelman, 1998). We described the posterior distributions for each parameter by their mean and 95% CI and assessed the strength of individual effects or the magnitude of difference between estimates based on the degree to which the 95% CI for the estimate did or did not overlap zero.

## 3 Results

We live-trapped 117,432 trap nights and captured 1,403 individual flying squirrels (692 females, 711 males) and 4,394 individual chipmunks (1,825 females, 2,569 males). Of these, we estimated ***φ*** for the 1,272 flying squirrels and 3,873 chipmunks that were captured before the final trapping occasion. Average site- and year-specific individual captures were 28.7 (range = 4–57) flying squirrels and 69.1 (range = 19–165) chipmunks. Site- and year-specific (2012–2019) counts of unmarked individuals ranged 0–12 for flying squirrels and 3–54 for chipmunks.

The most supported model of *p* for flying squirrel included the additive effects of a mean intercept and an individual-level random effect (Table 2; Table S1). Mean *p* of flying squirrels was 0.66 (95% CI: 0.51–0.73), but individual estimates varied substantially (***σ***_mH_ = 4.26, 95% CI: 3.19–4.96), ranged 0.25 (95% CI: 0.02–0.69) to 0.96 (95% CI: 0.76–0.99), and had a bi-modal posterior density with a dominant peak at approximately 0.45 and a smaller secondary peak at 0.81 (Figure 1). The most supported model of *p* for chipmunk included the additive effects of a mean intercept, a permanent behavioral trap response, and an individual-level random effect (Table 2; Table S1). Mean *p* of chipmunks was 0.37 (95% CI: 0.01–0.94), and the probability of recapture of individuals after encountering traps was 0.77 (95% CI: 0.74–0.80). Individual *p* of chipmunks varied substantially (***σ***_mH_ = 4.19, 95% CI: 2.9–4.96), ranged 0.21 (95% CI: 0.01–0.62) to 0.93 (95% CI: 0.61–0.99), and also had a bi-modal posterior density with a dominant peak at approximately 0.50 and a much smaller secondary peak at 0.83 (Figure 1).

**FIGURE 1.**
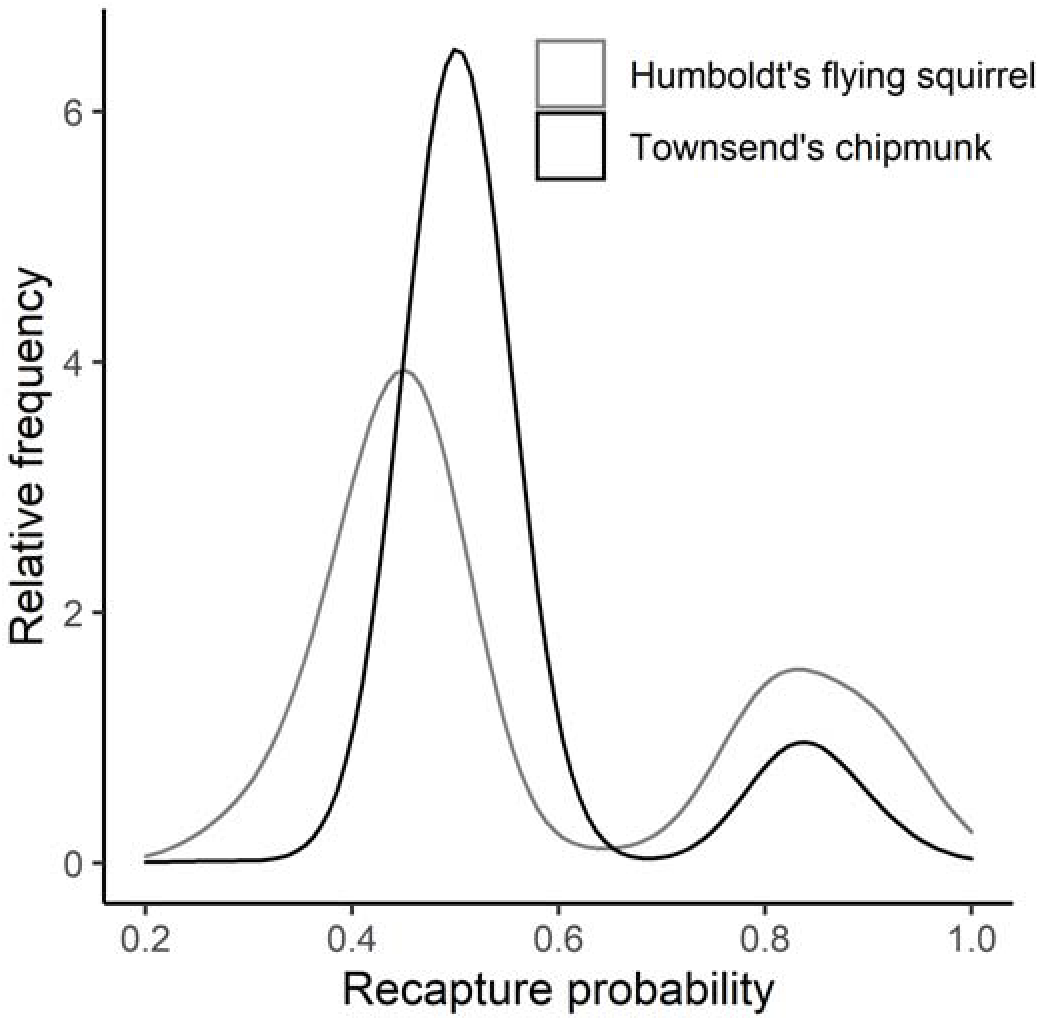
Relative frequencies of individual recapture probabilities for Humboldt’s flying squirrels (*Glaucomys oregonensis*) and Townsend’s chipmunks (*Neotamias townsendii*) estimated from models including an individual level random effect describing the observation process. The bimodal density plots are displayed with a gaussian kernel using a smoothing bandwidth of 0.05.

**TABLE 2.** Top three ranking models used to estimate recapture probability (*p*) of Humboldt’s flying squirrels (*Glaucomys oregonensis*; HFS) and Townsend’s chipmunks (*Neotamias townsendii*; TC) on nine late-successional forest sites in the H. J. Andrews Experimental Forest, 2011–2019. Column headings indicate the species, recapture probability model structure, the relative change in Watanabe–Akaike information criterion (ΔWAIC) from the top-ranking model, the lower 95% credible level for the relative change (Lower), the upper 95% credible interval for the relative change (Upper), and the model support weight (ω).

| Species | Model<sup>a</sup> | ΔWAIC | ΔWAIC 95% CI | | $\omega$ |
| --- | --- | --- | --- | --- | --- |
|  |  |  | Lower | Upper |  |
| HFS | mH | 0 | NA | NA | 1 |
|  | s | 32 | 12.195 | 51.805 | 0 |
|  | b + tRE | 48.43 | 26.907 | 69.953 | 0 |
|  | Null | 469.83 | 451.498 | 488.162 | 0 |
| TC | b + mH | 0 | NA | NA | 1 |
|  | mH | 91.24 | 71.168 | 111.312 | 0 |
|  | b + tRE | 449.28 | 428.737 | 469.823 | 0 |
|  | Null | 658.99 | 640.594 | 677.386 | 0 |

<sup>a</sup> Model structures are defined in Table 1.

Immigration rates varied among years for flying squirrels (***σ***_t_ = 2.88, 95% CI: 1.76–4.59), but less so for chipmunks (***σ***_t_ = 1.41, 95% CI: 0.83–2.55), and the species-specific pattern of temporal variation was similar for both sexes. Flying squirrel immigration rates for females and males were low (≤ 0.11) for both age-classes during six of eight study occasions (2013–2016, 2018, 2019), but were much higher (~ 0.22) for females and males of both age classes in 2012 and 2017 (Table S2). Similarly, chipmunk immigration rates were relatively low during six of eight study occasions (2012, 2014–2018), and were much higher for females and males during two occasions (2013, 2019; Table S2). Chipmunk immigration rates were higher relative to flying squirrel immigration rates during all occasions. We observed weak evidence that immigration rates of subadult and adult flying squirrels were lower than for juveniles, with < 10% of the coefficient 95% CI overlapping zero (*β*_Age_ = −0.13, 95% CI: −0.28–0.02). Female flying squirrels (*β*_Sex_ = −0.21, 95% CI: −0.36– −0.06) and chipmunks (*β*_Sex_ = −0.33, 95% CI: −0.41– −0.25) had lower immigration rates than males of those species respectively.

For flying squirrels, ***φ*** estimates varied among age, year, and sex (***σ***_a,s,t_ = 0.73, 95% CI: 0.46–1.06). For female flying squirrels, ***φ*** ranged 0.31 (95% CI: 0.12–0.55) to 0.71 (95% CI: 0.49–0.90) for juveniles, 0.43 (95% CI: 0.27–0.60) to 0.83 (95% CI: 0.69–0.94) for subadults, and 0.43 (95% CI: 0.32–0.55) to 0.84 (95% CI: 0.69–0.95) for adults (Figure 2). For male flying squirrels, ***φ*** ranged 0.29 (95% CI: 0.11–0.53) to 0.50 (95% CI: 0.29–0.73) for juveniles, 0.39 (95% CI: 0.19–0.61) to 0.70 (95% CI: 0.54–0.85) for subadults, and 0.37 (95% CI: 0.26–0.49) to 0.80 (95% CI: 0.61–0.94) for adults. Pairwise differences in ***φ*** and ***φ***_adjusted_ among flying squirrel age and sex classes were generally small and the 95% CIs for the pairwise differences broadly overlapped zero. We found weak evidence that female juvenile ***φ*** was lower relative to female subadult ***φ*** (*φ*_female juvenile_ – ***φ***_*female subadult*_ = −0.19, 95% CI. −0.41–0.03) and male juvenile ***φ*** was lower relative to both male subadult ***φ***((*φ_maie juveniie_* – *φ_maie subadult_* –0.20, 95% CI: −0.40–0.02) and male adult ***φ*** (*φ_male juvenile_* ~ *φ_male adult_* = −0.18, 95% CI: - 0.37–0.02). The rank order of ***φ*** among age classes was more variable among years for female flying squirrel’s relative to males, with all age classes represented as the minimum and maximum of the within-year estimates at least once during eight years (Figure 2). For male flying squirrels, juvenile ***φ*** was lowest among within-year estimates during all eight years, with the maximum of within-year estimates alternating between the subadult (maximum 5 of 8 years) and adult (maximum 3 of 8 years) age classes (Figure 2). Emigration adjustments varied temporally, ranged 0.01 (95% CI: 0.00–0.02) to 0.18 (95% CI: 0.10–0.27), and were substantial during the 2011–2012 and 2016–2017 intervals (Figure 3). The magnitude of differences in adjustments among age and sex classes were small (Table S3).

**FIGURE 2.**
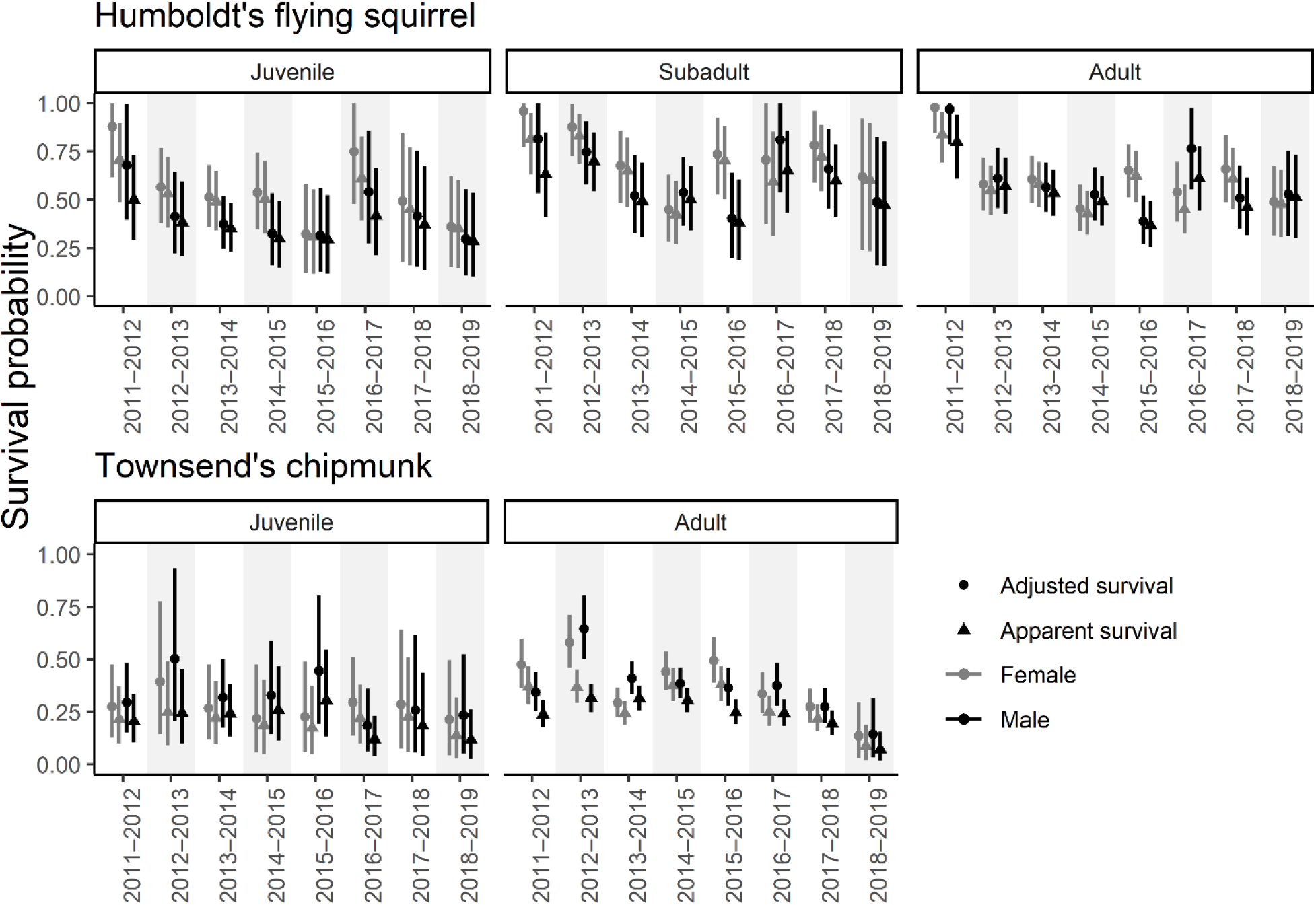
Estimates (mean and 95% credible interval) of annual apparent survival (***φ***, triangles) and adjusted survival (***φ***_adjusted_, circles) for female (grey) and male (black) Humboldt’s flying squirrels (*Glaucomys oregonensis*) and Townsend’s chipmunks (*Neotamias townsendii*), 2011–2019, in the H. J. Andrews Experimental Forest in Oregon.

**FIGURE 3.**
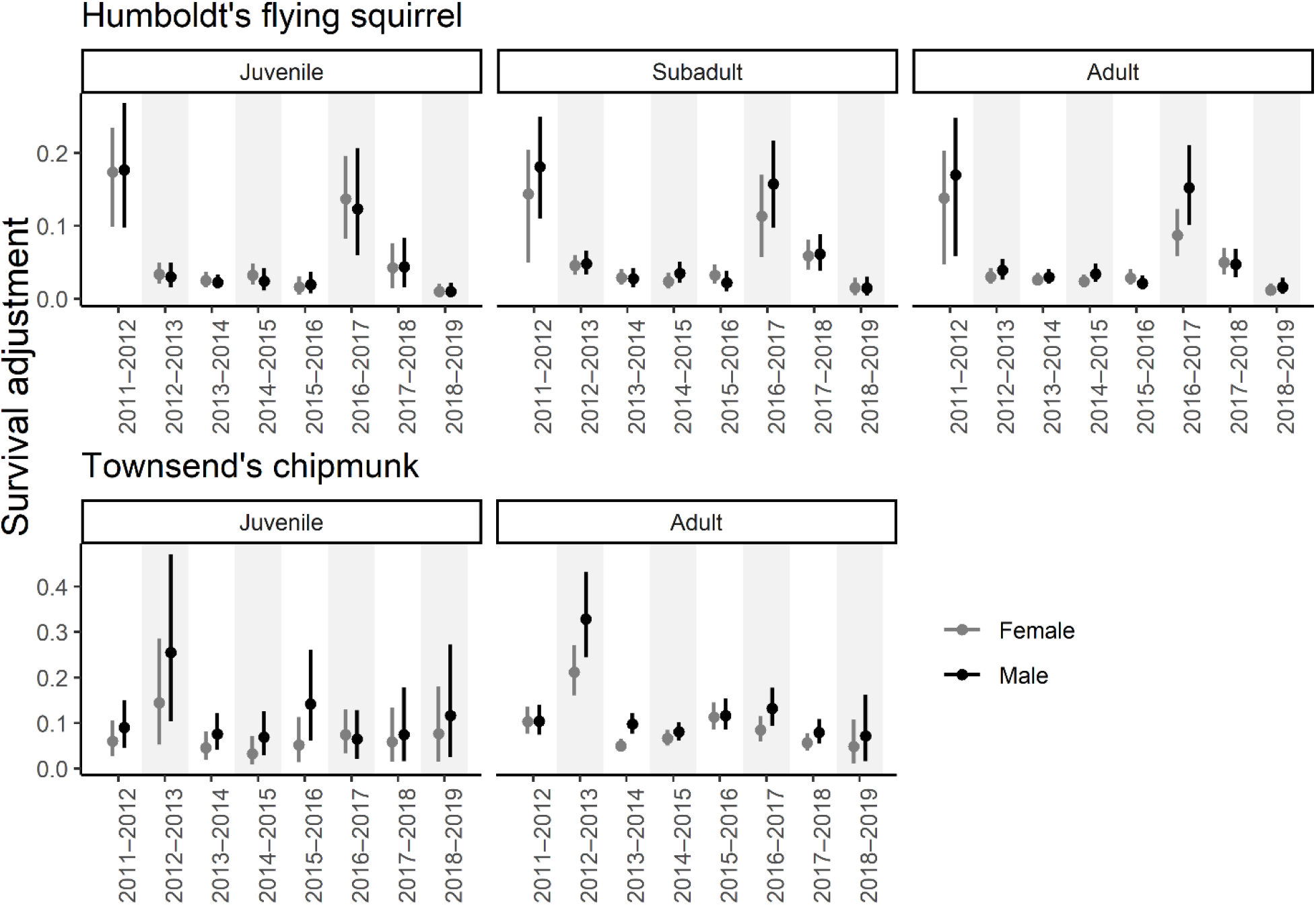
Age- and sex-specific apparent survival emigration-adjustment (Δ_*survival adjustment*_ = φ_adjusted_ – φ) estimates (mean and 95% credible intervals) for Humboldt’s flying squirrels (*Glaucomys oregonensis*) and Townsend’s chipmunks (*Neotamias townsendii*) captured 2011–2019 on the H. J. Andrews Experimental Forest in Oregon.

For chipmunks, annual ***φ*** varied among years, but differed much less among age and sex relative to flying squirrels (***σ***_a,s,t_ = 0.69, 95% CI: 0.39–0.1.13). Annual ***φ*** ranged 0.13 (95% CI: 0.03–0.32) to 0.25 (95% CI: 0.09–0.50) for juvenile females and ranged 0.09 (95% CI: 0.02–0.19) to 0.38 (95% CI: 0.30–0.46) for adult females (Figure 2). For males, annual ***φ*** ranged 0.11 (95% CI: 0.03–0.26) to 0.30 (95% CI: 0.13–0.55) for juveniles and 0.07 (95% CI: 0.02–0.16) to 0.31 (95% CI: 0.26–0.37) for adults (Figure 2). Pairwise differences in ***φ*** and ***φ***_adjusted_ among each chipmunk sex and age class were small and the 95% CI for each difference broadly overlapped zero. For both sexes, the rank order of juvenile ***φ*** was lower than adult survival during most years (female: 7 of 8 years, male: 6 of 8 years). Emigration adjustments ranged 0.03 (95% CI: 0.01–0.07) to 0.33 (95% CI: 0.24–0.43), varied temporally for both sexes, and were substantial during the 2012–2013 interval (Figure 3). Emigration adjustments were larger in magnitude for males relative to females during all years (Table S3).

Covariate posterior distributions were similar for both sets of priors (Table 3, Figure S2). Visual inspection of trace plots and estimates of the Brooks–Gelman–Rubin convergence diagnostic indicated convergence was obtained for all monitored parameter estimates 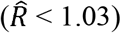. Bayesian p-values estimated from the posterior predictive checks were 0.53 for flying squirrels and 0.44 for chipmunks, indicating adequate fit for all models and suggesting that both candidate models generated data consistent with the observed data.

**TABLE 3.**
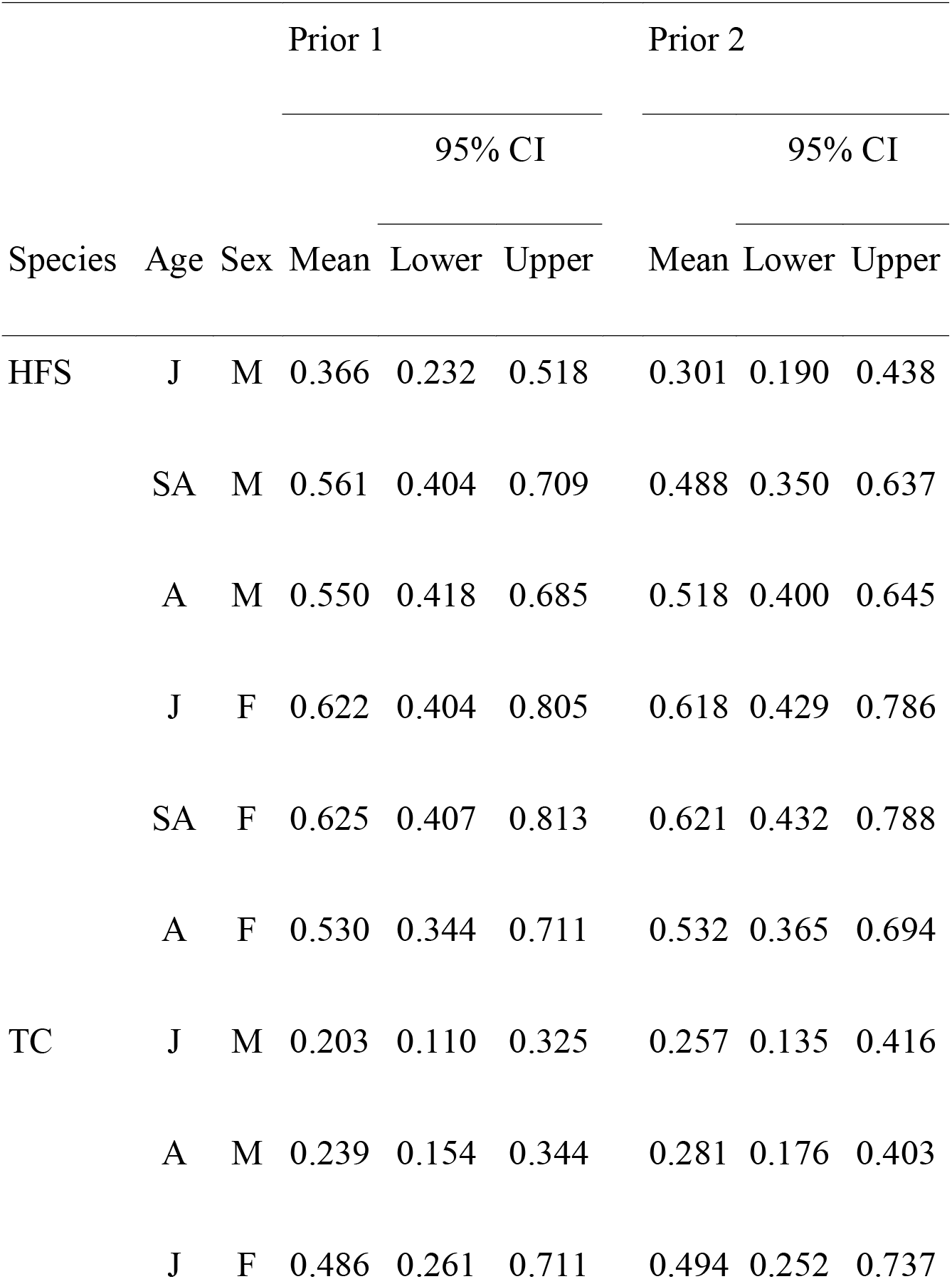
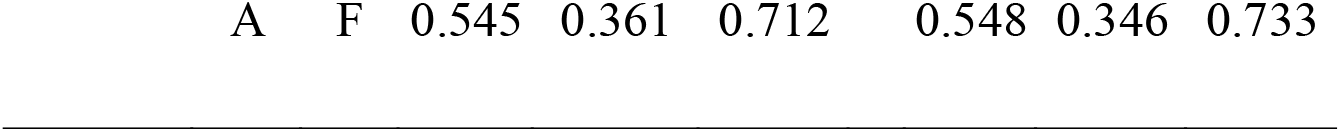
Time-invariant age- and sex-specific estimates of mean apparent annual survival (***φ***; real scale) and associated 95% credible intervals (lower: 2.5%, upper: 97.5%) for Humboldt’s flying squirrels (*Glaucomys oregonensis*; HFS) and Townsend’s chipmunks (*Neotamias townsendii*; TC) captured 2011–2019 on the H. J. Andrews Experimental Forest in Oregon. Super columns ‘Prior 1’ and ‘Prior 2’ refer to estimates obtained using two covariate prior sets, strong differences would indicate model sensitivity to prior selection. Age and sex categories include juvenile (J), subadult (SA), adult (A), male (M), and female (F).

|  |  |  | Prior 1 |  |  | Prior 2 |  |  |
| --- | --- | --- | --- | --- | --- | --- | --- | --- |
|  |  |  | 95% CI |  |  | 95% CI |  |  |
| Species | Age | Sex | Mean | Lower | Upper | Mean | Lower | Upper |
| HFS | J | M | 0.366 | 0.232 | 0.518 | 0.301 | 0.190 | 0.438 |
|  | SA | M | 0.561 | 0.404 | 0.709 | 0.488 | 0.350 | 0.637 |
|  | A | M | 0.550 | 0.418 | 0.685 | 0.518 | 0.400 | 0.645 |
|  | J | F | 0.622 | 0.404 | 0.805 | 0.618 | 0.429 | 0.786 |
|  | SA | F | 0.625 | 0.407 | 0.813 | 0.621 | 0.432 | 0.788 |
|  | A | F | 0.530 | 0.344 | 0.711 | 0.532 | 0.365 | 0.694 |
| TC | J | M | 0.203 | 0.110 | 0.325 | 0.257 | 0.135 | 0.416 |
|  | A | M | 0.239 | 0.154 | 0.344 | 0.281 | 0.176 | 0.403 |
|  | J | F | 0.486 | 0.261 | 0.711 | 0.494 | 0.252 | 0.737 |

|  |  |  |  |  |  |  |  |
| --- | --- | --- | --- | --- | --- | --- | --- |
| A | F | 0.545 | 0.361 | 0.712 | 0.548 | 0.346 | 0.733 |

## 4 Discussion

Our analysis provided estimates of age-, sex-, and time-specific apparent annual survival and emigration-adjusted survival for two small mammal species captured in a late successional forest 2011–2019, and reinforces the need to adjust for emigration in estimates of apparent survival. Here we also present analytical methods to parse confounded survival and movement probabilities to reduce bias in empirical apparent annual survival estimates relative to emigration-adjusted survival. By reducing bias associated with emigration, we revealed that juvenile flying squirrel survival may be much higher than expected as our estimates are among the highest observed for any small mammal species (Table S4; Kraus et al., 2005). Indeed, our emigration-adjusted survival for juvenile flying squirrels exceeded reported adult apparent annual survival estimates for many other species (Schaub & Vaterlaus-Schelegel, 2001), including adult chipmunks in our study. Our approach also uncovered temporal variation in apparent survival biases that are likely applicable to a wide range of species and ecosystems.

Our findings were consistent with previous studies that demonstrated sometimes-substantial negative biases of apparent survival relative to true survival, due to confounding among survival and emigration probabilities (Lebreton et al., 1992; Schaub & Royle, 2014). Beyond that widely-recognized phenomenon (e.g., Schaub et al., 2004; Schaub & Royle, 2014), we demonstrated that levels of bias in apparent survival can vary substantially over time due to temporal variation in site-fidelity rates. For both species examined here, the bias induced by confounding of emigration and survival probabilities was not consistent among years. In general, we observed consistent agreement between apparent survival and adjusted survival estimates for both species, sexes, and all age-classes. However, during some years, the bias for one or more age- and sex-specific classes was >1 order of magnitude larger than the bias in other years, and during those years, inferences for temporal variation in apparent survival and adjusted survival differed. For example, our estimates of apparent annual survival for juvenile and subadult male flying squirrels during 2016–2017 and 2017–2018 are similar, whereas mean emigration-adjusted survival is much higher during 2016–2017 relative to 2017–2018. This is concerning for unadjusted apparent survival estimates, especially for short-term studies that cannot differentiate between years when apparent survival is a suitable estimator for survival and years when movement probabilities are important and influential confounders. Moreover, differences in survival probabilities or movement behavior among age or sex classes could further confound estimation of either quantity individually (Schaub & Royle, 2014).

For flying squirrels, our estimates of adult apparent annual survival and adjusted survival were intermediate to previously-reported estimates that ranged 0.32–0.68 (Gomez, 2005; Lehmkuhl et al., 2006; Ransome & Sullivan, 2002), including those reported by Weldy et al. (2020) for data collected on these sites 2011–2016. In that analysis, temporal variation in apparent annual survival was not supported by model selection criteria, consistent with the findings of Lehmkuhl et al. (2006). But, in this analysis, we focused on estimates of age- and sex-specific survival and estimated substantial temporal variation in apparent annual survival, especially for the juvenile and subadult age-classes. In the expanded time series of mark-recapture data used here, our estimates of apparent annual survival during 2012–2016 were similar to those reported by Weldy et al. (2020) and showed little temporal variation, especially for the adult age class. However, during 2016–2019, we observed a peak and subsequent decline in apparent survival and emigration-adjusted survival. For chipmunks, our estimates of apparent annual survival and adjusted survival were similar to conspecifics and congenerics (Schulte-Hostedde et al., 2002; Weldy et al., 2020). But we found less evidence for temporal variation in apparent annual survival, and temporal variation in adjusted survival did not match previously reported patterns (Weldy et al. 2020).

Consistent with our hypotheses, juvenile flying squirrel survival was lower relative to subadults or adults, and male survival was lower relative to females for all three age classes, but the magnitude of that difference diminished with age. We also expected lower survival in juvenile chipmunks, but we found little difference in age- or sex-specific survival. For both species, however, temporal variation of survival within age and sex classes was larger than variation among age and sex classes, highlighting the importance of long-term studies to understand variation of demographic traits.

The importance of estimating age-specific survival while correcting for emigration is demonstrated by contrasting findings of this study with those of previous research in this system. Weldy et al. (2020) observed a negative association between apparent annual survival and recruitment rate for chipmunks, where low apparent survival was coupled with high recruitment and low recruitment was coupled with high survival. If age-specific survival were a primary driver of this observation, we expected to observe relatively stable survival of adult chipmunks while juvenile survival varied. Instead, we conclude that the low survival estimates coupled with high recruitment were associated with individual movement. Recruitment was large because individuals moved into study populations, while survival was low because marked individuals left those populations.

Immigration rates of flying squirrels were generally low, except in 2012 and 2017 when estimated immigration rates were more than 2-fold higher. For chipmunks, in comparison, immigration rates were more variable and much higher overall. For example, the lowest estimates of chipmunk immigration rates were nearly equivalent to the two peak flying squirrel immigration estimates. Taken together, these estimates demonstrate different temporal patterns of immigration, and consequently temporal variation in the influence of emigration on estimates of apparent survival. During most years, immigration is likely an unimportant driver of flying squirrel population dynamics, whereas it is likely consistently influential to chipmunk population dynamics.

We were concerned that age- or sex-specific variation in site-fidelity rates would bias inferences about differences in survival. We chose to use a sub-model extension to the CJS model framework that estimated age-, sex-, and time-specific immigration probabilities, which we used to derive estimates of site-fidelity rates and adjusted survival. This approach is similar to others that integrate data to estimate adjusted survival (Abadi et al., 2010), which typically parameterize immigration as a Poisson-distributed rate or count (Schaub & Fletcher, 2015). Early integrated analyses suggested that one crucial assumption of integrated models is that the datasets to be integrated are independent (Besbeas et al., 2002; Schaub & Abadi, 2011); violation of that assumption is thought to result in overestimates of precision (Lebreton et al., 1992). However, Weegman et al. (2020) found no effects on parameter bias or precision from integrated population models fit to simulated data with complete overlap.

Employing the emigration correction on survival estimates by our method requires carefully considering whether source or sink habitats exist within the study area. Our immigration sub-model extension was contingent on the assumption that emigration rates at time *t* are equivalent to immigration rates at time *t*+1 and that the site-fidelity rates were complement to emigration rates. This is equivalent to the assumption that our study species are moving through the study system randomly in an even flow within a year. We felt that this assumption was met on our study-sites, which were randomly placed within a large, continuous, old late-successional forest where site edges did not reflect biological edges. Further, Carey (1995) suggested that there was no evidence that the densities of flying squirrels and chipmunks were misleading indicators of habitat suitability, and Weldy et al. (2019) found no evidence for marginal or sink habitat on our study sites. Presence of sink habitats would have indicated that individuals were more likely to emigrate from or immigrate to specific sites. In that case, apparent annual survival estimates would be biased low, and violation of our random and even movement adjustment assumption would have caused our adjustment to apparent annual survival to overcompensate for the negative biases caused by movement on some sites.

Our study demonstrates a novel approach to gaining insight into links between movement and survival for two species of small mammals. The study of small mammal movement ecology often lags behind that of larger species, leading to knowledge gaps such as how movement of small mammals influences ecosystem health through the dispersal of hypogeous fungi spores and plant seeds (Trappe et al., 2009). However, while our methods are appropriate given the homogenous nature of our study area, the necessary assumptions might not hold in other systems with more fragmentation, variable habitat quality, or potential for source-sink dynamics among sites. Future studies could continue to explore the movement ecology of small mammals by incorporating study designs suitable for directly estimating movement parameters (i.e., multistate mark-recapture or telemetry) and evaluating the effects of spatiotemporal predictors on movement probabilities, or by linking temporal variation of movement rates to studies of population cycling (e.g., Fryxell et al., 1998; Weldy et al., 2019).

## 5 Acknowledgements

Funding for small-mammal trapping and analysis was provided by the USDA Forest Service, Pacific Northwest Research Station and conducted on federal lands with approval from the USDA Forest Service. Facilities were provided by the H. J. Andrews Experimental Forest and Long-term Ecological Research (LTER) program, administered cooperatively by the USDA Forest Service Pacific Northwest Research Station, Oregon State University, and the Willamette National Forest. This material is based upon work supported by the National Science Foundation under the LTER Grants: LTER8 DEB-2025755 (2020-2026) and LTER7 DEB-1440409 (2012-2020). We thank T. M. Manning and T. M. Wilson for their guidance, support, and logistical management. For field work, we thank N. B. Alexander, D. A. Arnold, A. M. Bartelt, A. Bies, S. C. Bishir, N. A. Bromen, L. L. Carver, M. J. Cokeley, C. R. Gray, L. Hodnett, L. K. Howard, A. C. Hsiung, J. Hurd, C. J. Hutton, D. K. Jacobsma, P. C. Kannor, B. E. Kerfoot, M. A. Linnell, T. J. Mayer, L. S. Millward, B. Nahorney, T. Newton, H. M. Oswald, S. M. Pack, K. A. Ray, A. Rutherford, D. C. Tange, K. Van Neste, S. E. Ward, W. Watson, and J. M. Winiarski. This publication represents the views of the authors, and any use of trade, firm, or product names is for descriptive purposes only and does not imply endorsement by the United States Government.

## Author Contributions

MW conceived the ideas, collected data, led manuscript writing, designed methodology and analyzed data. All authors contributed to the drafts, interpretation of results, and gave final approval for publication.

## Data Availability Statement

Data used to estimate apparent annual survival and immigration for flying squirrels and chipmunks are available from figshare: https://figshare.com/projects/Small_Mammal_Age-Specific_Survival/119784. R and JAGS code to recreate the analyses and figures is available on GitHub https://github.com/MJWeldy/SM_MAMM_AGE_SURVIVAL.

## Supporting Information

**FIGURE S1.**
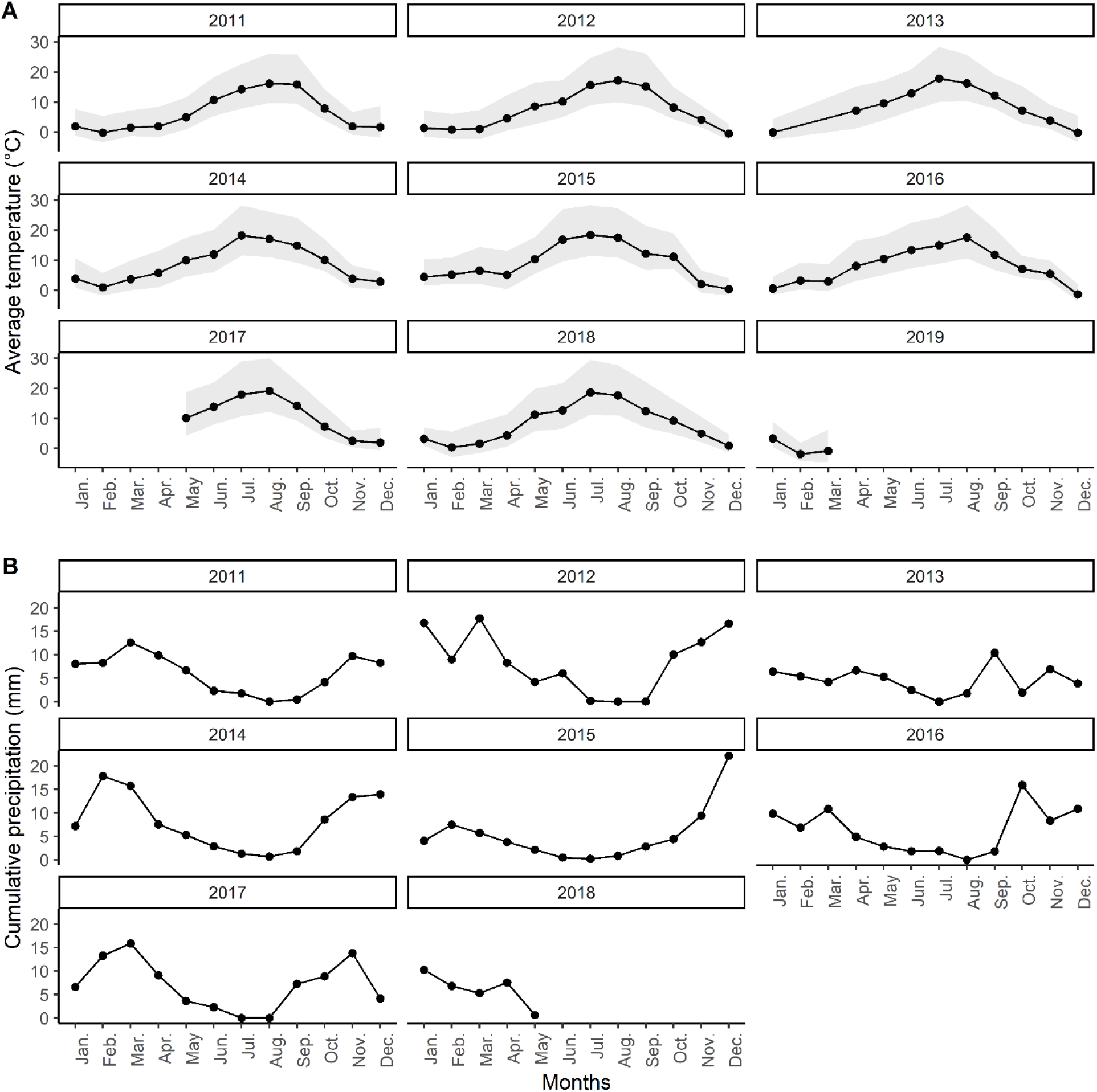
A) Average monthly temperatures (°C; April 2011 to March 2019) and B) cumulative monthly precipitation (mm; April 2011 to September 2018) recorded at the central meteorological station (1,020 m) weather station in the H. J. Andrews Experimental Forest. The grey color band around average monthly temperature values indicates the range (minimum to maximum) of temperatures recorded within the month.

**FIGURE S2.**
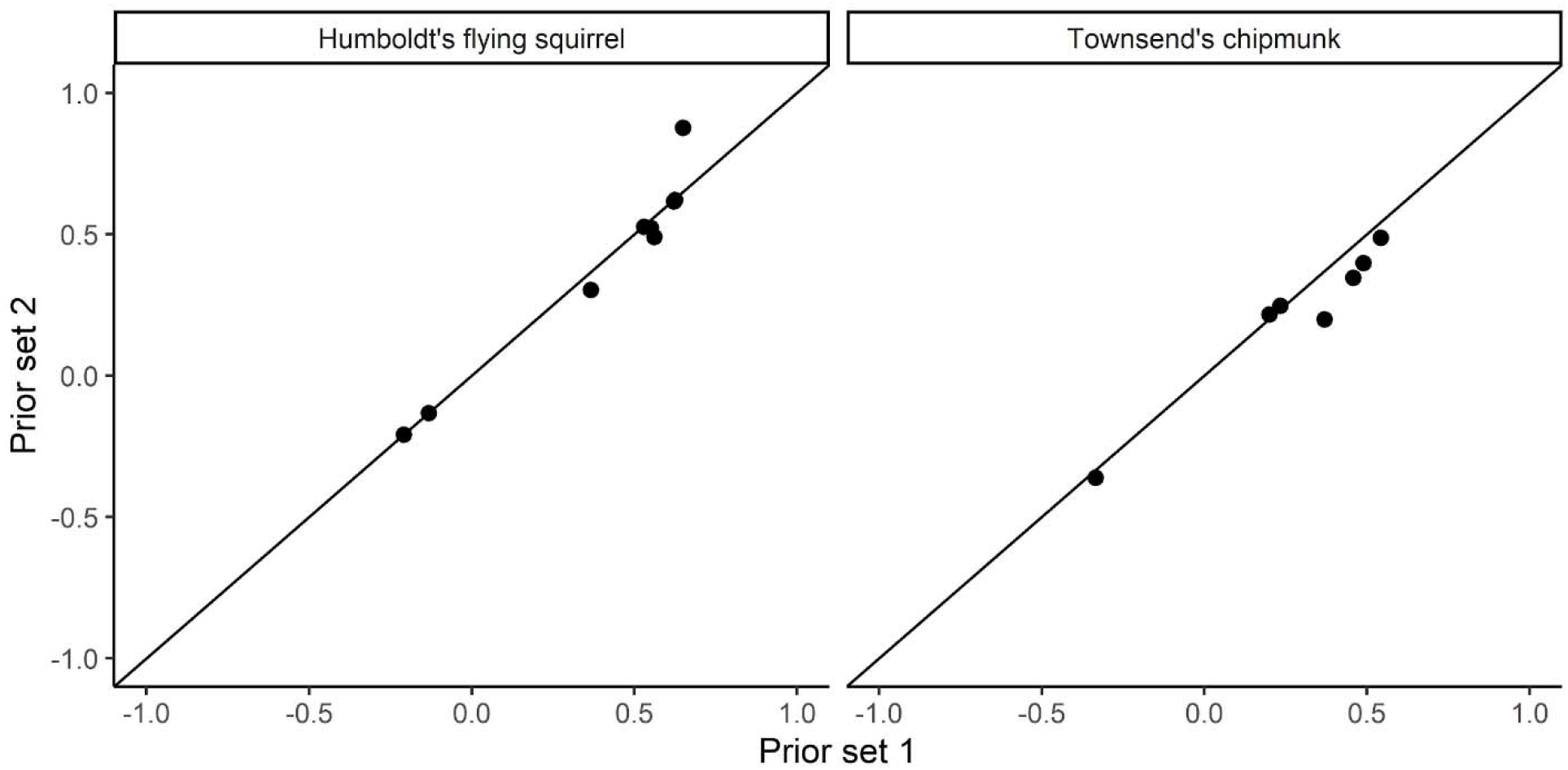
Prior sensitivity plot for uninformative prior sets 1 and 2 used to estimate apparent annual survival, recapture probability, and immigration rates of Humboldt’s flying squirrels (*Glaucomys oregonensis*) and Townsend’s chipmunks (*Neotamias townsendii*) captured during 2011–2019 in the H. J. Andrews Experimental Forest in Oregon.

**TABLE S1.**
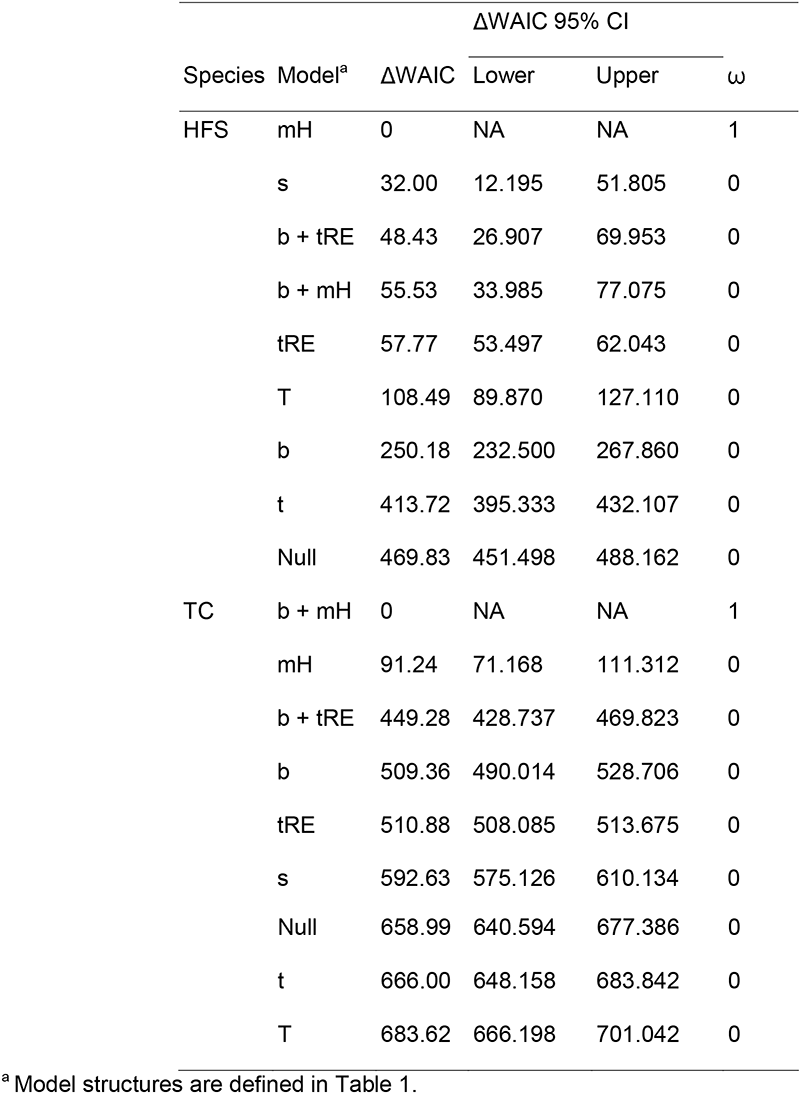
Model selection results used to estimate recapture probability (*p*) of Humboldt’s flying squirrels (*Glaucomys oregonensis*; HFS) and Townsend’s chipmunks (*Neotamias townsendii*; TC) on nine late-successional forest sites in the H. J. Andrews Experimental Forest 2011–2019. Column headings indicate the species, recapture probability models structure, the relative change in Watanabe–Akaike information criterion (ΔWAIC) from the top-ranking model, the lower 95% credible level for the relative change (Lower), the upper 95% credible interval for the relative change (Upper), and the model support weight (ω).

| Species | Model<sup>a</sup> | ΔWAIC | ΔWAIC 95% CI | | $\omega$ |
| --- | --- | --- | --- | --- | --- |
|  |  |  | Lower | Upper |  |
| HFS | mH | 0 | NA | NA | 1 |
|  | s | 32.00 | 12.195 | 51.805 | 0 |
|  | b + tRE | 48.43 | 26.907 | 69.953 | 0 |
|  | b + mH | 55.53 | 33.985 | 77.075 | 0 |
|  | tRE | 57.77 | 53.497 | 62.043 | 0 |
|  | T | 108.49 | 89.870 | 127.110 | 0 |
|  | b | 250.18 | 232.500 | 267.860 | 0 |
|  | t | 413.72 | 395.333 | 432.107 | 0 |
|  | Null | 469.83 | 451.498 | 488.162 | 0 |
| TC | b + mH | 0 | NA | NA | 1 |
|  | mH | 91.24 | 71.168 | 111.312 | 0 |
|  | b + tRE | 449.28 | 428.737 | 469.823 | 0 |
|  | b | 509.36 | 490.014 | 528.706 | 0 |
|  | tRE | 510.88 | 508.085 | 513.675 | 0 |
|  | s | 592.63 | 575.126 | 610.134 | 0 |

|  |  |  |  |  |
| --- | --- | --- | --- | --- |
| Null | 658.99 | 640.594 | 677.386 | 0 |
| t | 666.00 | 648.158 | 683.842 | 0 |
| T | 683.62 | 666.198 | 701.042 | 0 |

<sup>a</sup> Model structures are defined in Table 1.

**TABLE S2.**
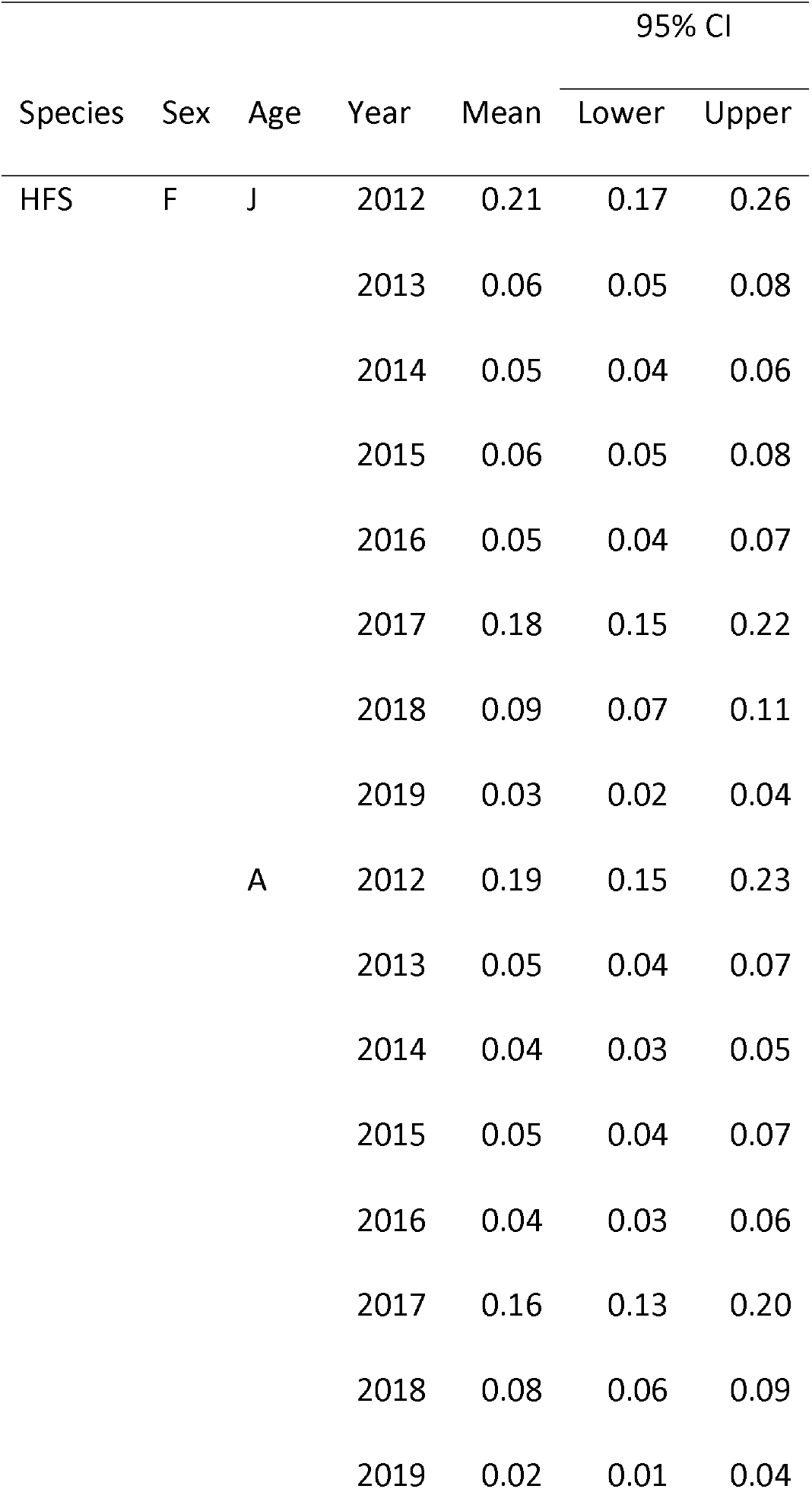
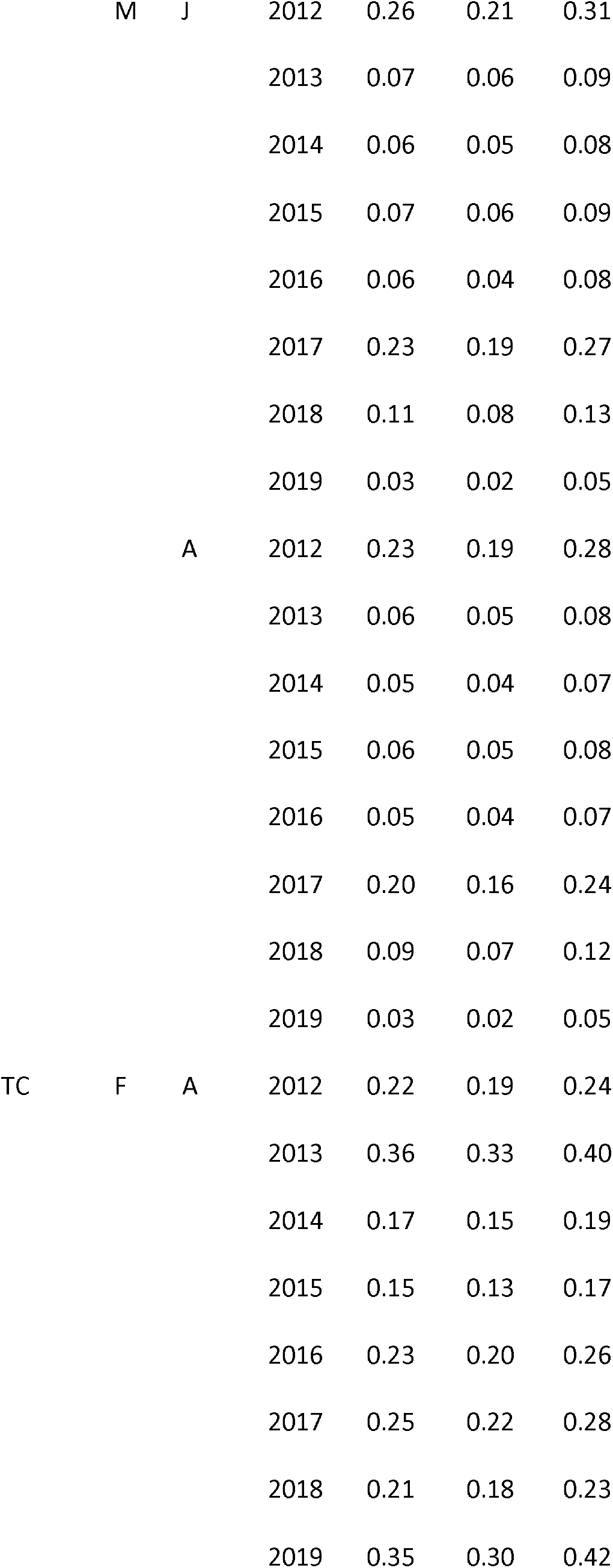
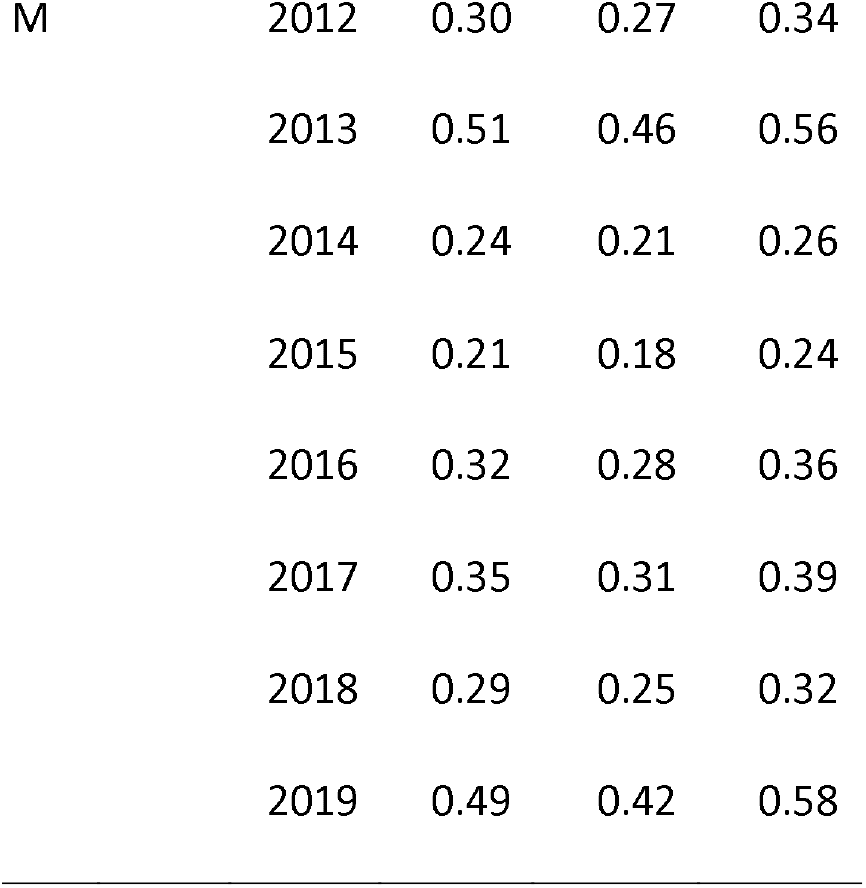
Age- and sex-specific estimates of immigration rates and associated 95% credible intervals (lower: 2.5%, upper: 97.5%) for Humboldt’s flying squirrels (*Glaucomys oregonensis*; HFS) and Townsend’s chipmunks (*Neotamias townsendii*; TC) captured 2011–2019 on the H. J. Andrews Experimental Forest in Oregon. Age and sex categories include juvenile (J), subadult (SA), adult (A), male (M), and female (F).

**TABLE S3.**
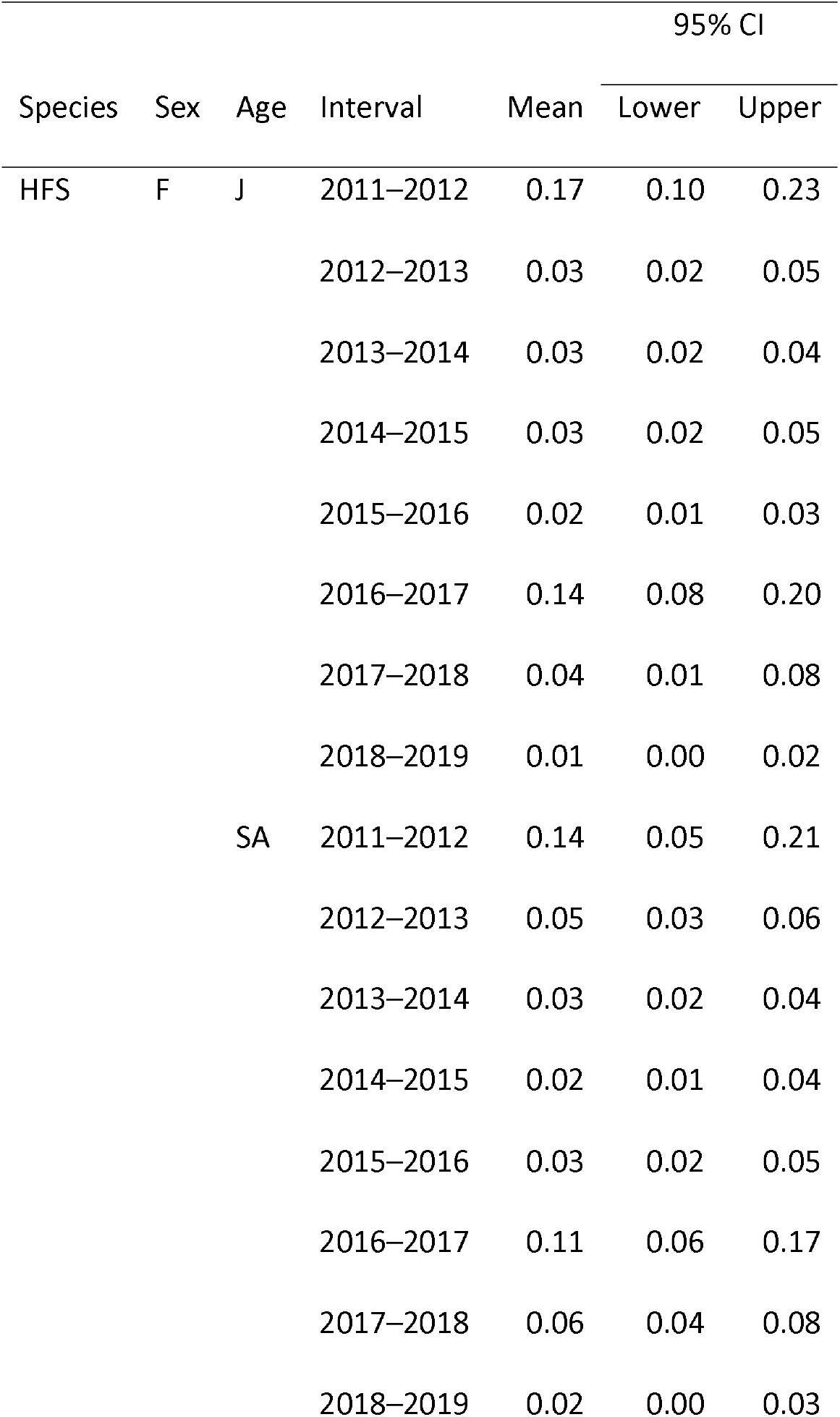
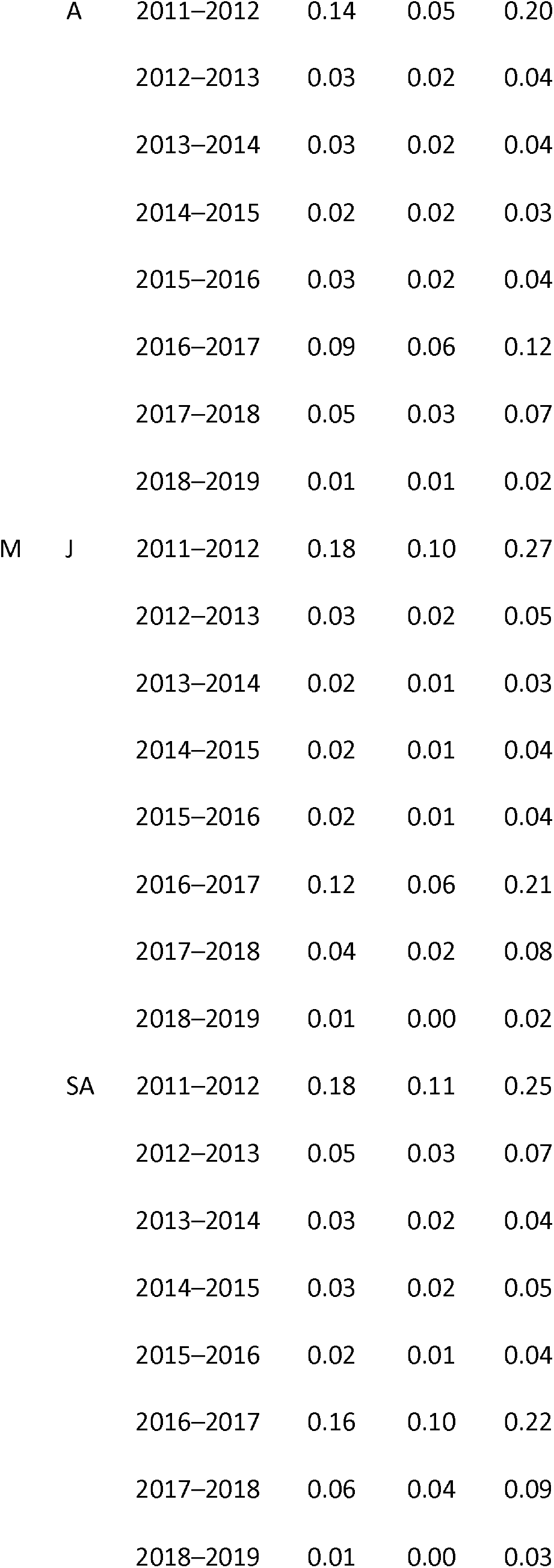
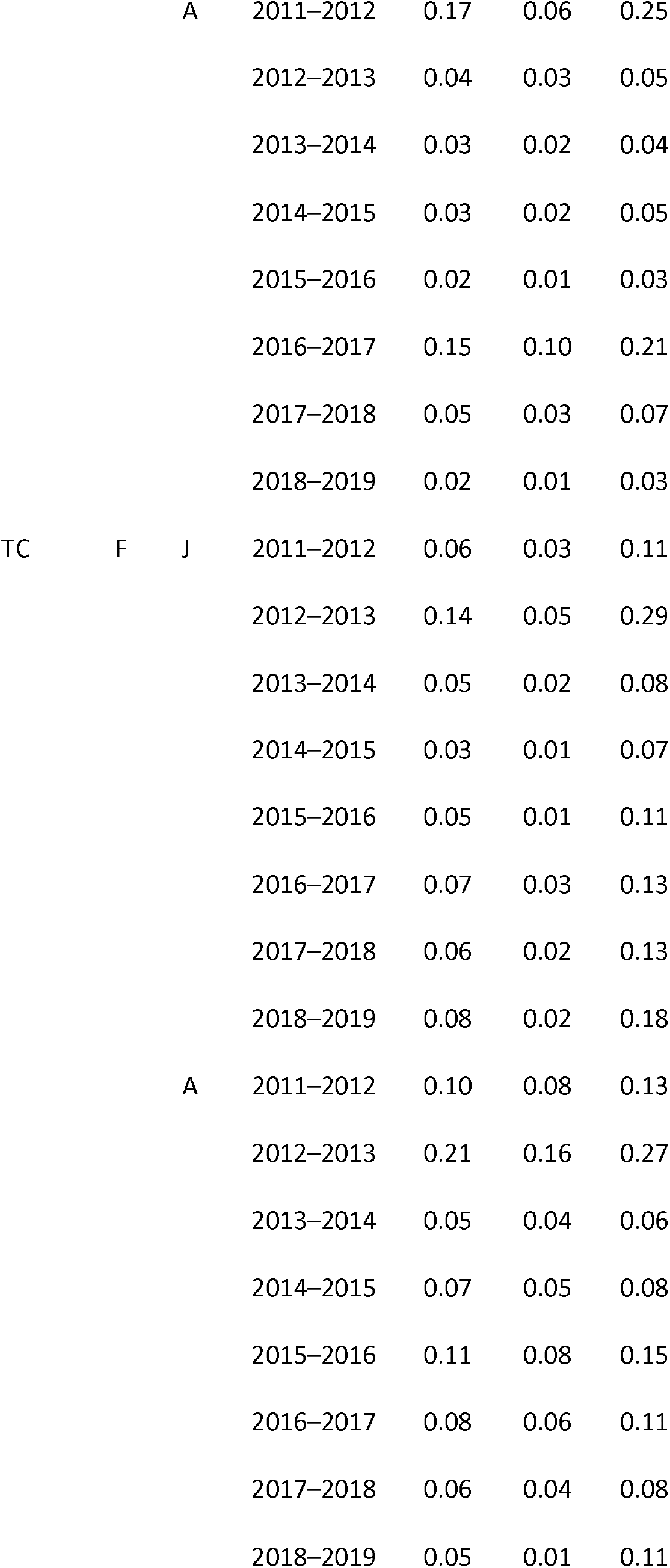
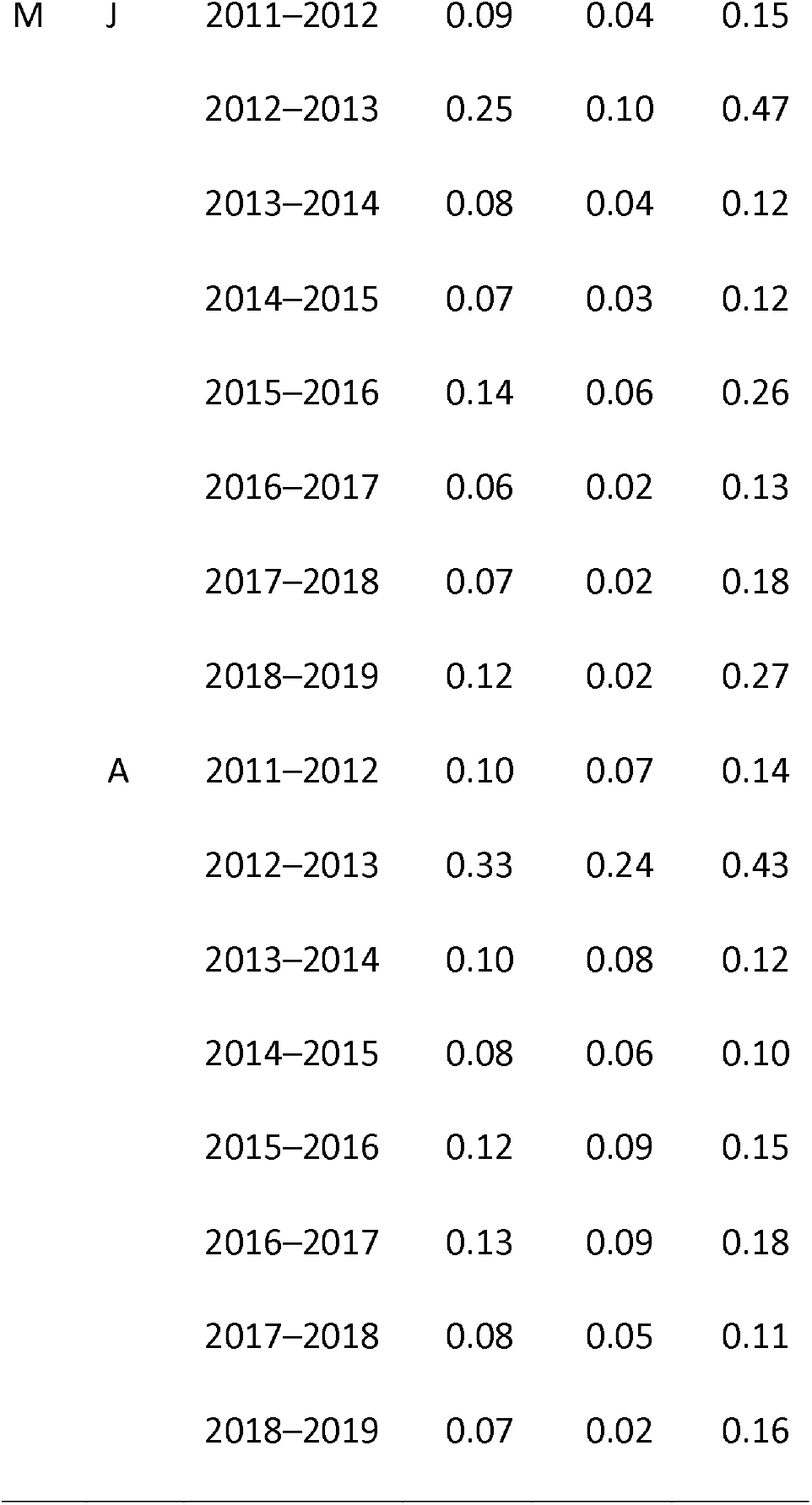
Age- and sex-specific apparent survival emigration-adjustment estimates and associated 95% credible intervals (lower: 2.5%, upper: 97.5%) for Humboldt’s flying squirrels (*Glaucomys oregonensis*; HFS) and Townsend’s chipmunks (*Neotamias townsendii*; TC) captured 2011–2019 on the H. J. Andrews Experimental Forest in Oregon. Age and sex categories include juvenile (J), subadult (SA), adult (A), male (M), and female (F).

**TABLE S4.** Ratios of adult to juvenile survival, including apparent annual survival (***φ***) and adjusted annual survival (***φ***_adjusted_), for male (M) and female (F) Humboldt’s flying squirrels (*Glaucomys oregonensis*; HFS) and Townsend’s chipmunks (*Neotamias townsendii*; TC) captured during 2011–2019 on the H. J. Andrews Experimental Forest in Oregon.

| Species | Estimate | Sex | Mean | Min | Max |
| --- | --- | --- | --- | --- | --- |
| HFS | $\phi$ | F | 1.22 | 0.74 | 2.02 |
|  |  | M | 1.49 | 1.24 | 1.78 |
| | $\phi_{\text{adjusted}}$ | F | 1.19 | 0.72 | 2.01 |
|  |  | M | 1.46 | 1.22 | 1.78 |
| TC | $\phi$ | F | 1.40 | 0.62 | 2.18 |
|  |  | M | 1.18 | 0.61 | 2.02 |
| | $\phi_{\text{adjusted}}$ | F | 1.40 | 0.62 | 2.18 |
|  |  | M | 1.18 | 0.61 | 2.03 |

